# Interaction Between HCN and Slack Channels Regulates mPFC Pyramidal Cell Excitability and Working Memory

**DOI:** 10.1101/2023.03.04.529157

**Authors:** Jing Wu, Lynda El-Hassar, Dibyadeep Datta, Merrilee Thomas, Yalan Zhang, P. Jenkins David, Nicholas J. DeLuca, Manavi Chatterjee, Valentin K. Gribkoff, Amy F.T. Arnsten, Leonard K. Kaczmarek

**Author notes:** **Corresponding authors:** Leonard K Kaczmarek, Department of Pharmacology, Yale School of Medicine, New Haven, Connecticut 06520; Department of Cellular and Molecular Physiology, Yale School of Medicine, New Haven, Connecticut 06520;. These authors contributed equally to this work (Jing Wu, Lynda El-Hassar).

## Abstract

The ability of monkeys and rats to carry out spatial working memory tasks has been shown to depend on the persistent firing of pyramidal cells in the prefrontal cortex (PFC), arising from recurrent excitatory connections on dendritic spines. These spines express hyperpolarization-activated cyclic nucleotide-gated (HCN) channels whose open state is increased by cAMP signaling, and which markedly alter PFC network connectivity and neuronal firing. In traditional neural circuits, activation of these non-selective cation channels leads to neuronal depolarization and increased firing rate. Paradoxically, cAMP activation of HCN channels in PFC pyramidal cells reduces working memory-related neuronal firing. This suggests that activation of HCN channels may hyperpolarize rather than depolarize these neurons. The current study tested the hypothesis that Na^+^ influx through HCN channels activates Na^+^-activated K^+^ (K_Na_ or Slack) channels to hyperpolarize the membrane. We have found that HCN and Slack K_Na_ channels co-immunoprecipitate in cortical extracts and that, by immunoelectron microscopy, they colocalize at postsynaptic spines of PFC pyramidal neurons. A specific blocker of HCN channels, ZD7288, reduces K_Na_ current in pyramidal cells that express both HCN and Slack channels, indicating that blockade of HCN channels reduced K^+^ current indirectly by lowering Na^+^ influx. In contrast, ZD7288 has no effect on K_Na_ currents in an HEK cell line stably expressing this Slack channels but no HCN channels, demonstrating that ZD7288 does not block Slack channels directly. Activation of HCN channels by cAMP in a cell line expressing a Ca^2+^ reporter results in elevation of cytoplasmic Ca^2+^, but the effect of cAMP is completely reversed if the HCN channels are co-expressed with Slack channels. Finally, we have used a novel pharmacological blocker of Slack channels to show that inhibition of either Slack or HCN channels in rat PFC improves working memory performance, and that the actions of Slack and HCN channel blockers occlude each other in the memory task. Our results suggest that the regulation of working memory by HCN channels in PFC pyramidal neurons is mediated by an HCN-Slack channel complex that links activation HCN channels to suppression of neuronal excitability.

## Introduction

Neuronal networks of the prefrontal cortex (PFC) subserve working memory, the ability to retain short-term neuronal information in the absence of sensory stimulation. This aspect of mental representation has been extensively studied in monkeys and rats using spatial working memory tasks, where the persistent firing of layer III neurons across the delay period is considered the cellular basis for spatial working memory (Funahashi et al., 1989). Persistent neuronal firing arises from extensive recurrent excitatory connections, including on dendritic spines in layer III of macaque dorsolateral PFC (dlPFC) (Goldman-Rakic, 1995). Persistent firing has also been found in rat medial PFC (mPFC) (Devilbiss et al., 2017; Yang et al., 2018), although with a shorter duration than in primates. Disruption of PFC pyramidal cell firing during stress or mental illness leads to working memory deficits and to inappropriate behavior (Nook et al., 2018; Arnsten et al., 2021). One mechanism by which stress can disrupt working memory performance is through excessive release of catecholamines, which activate cAMP signaling to influence the activity of ion channels that alter PFC pyramidal cell firing (Arnsten, 2015). cAMP signaling can increase the open state of hyperpolarization-activated cyclic nucleotide-gated (HCN) channels, non-selective cation channels that flux Na^+^ into the neuron (Chen et al., 2001). Although HCN channels flux an inward, excitatory current, their ultimate influences are often complex, e.g. altering membrane resistance and neuronal excitability (He et al., 2014; Kim et al., 2018). In macaque dlPFC, local increases in cAMP signaling dlPFC have been shown to reduce task-related neuronal firing and impair working memory performance by opening HCN channels (Wang et al., 2007), while low dose blockade of HCN channels enhances firing (Wang et al., 2007). Similarly, blockade of HCN channels in rat mPFC can prevent stress-induced working memory deficits (Gamo et al., 2015). Thus, understanding HCN channel mechanisms is important for revealing the etiology of cognitive deficits.

An important clue regarding HCN channel functional contributions comes from their differing subcellular locations within neurons in diverse circuits. The most common location for HCN channels on pyramidal cells is on the shaft of their distal apical dendrites, e.g. in macaque primary visual cortex (Yang et al., 2018), rodent hippocampus (Nolan et al., 2004), and deep layers of rodent cortex (Yang et al., 2018), including distal dendrites of layer V mPFC (Lörincz et al., 2002; Notomi and Shigemoto, 2004; Day et al., 2005). In layer III dlPFC, however, HCN channels are concentrated on dendritic spines near glutamatergic-like synapses (Paspalas et al., 2013b), where they are positioned to gate recurrent excitatory inputs needed for working memory (Wang et al., 2007). In dlPFC, HCN channels on dlPFC spines are co-localized with multiple cAMP signaling proteins, while those on dlPFC distal dendritic shafts are not (Paspalas et al., 2013b), suggesting differing regulation and functions. Indeed, in contrast to some neurons where HCN channel opening is excitatory (Thuault et al., 2013), cAMP opening of HCN channels in the dlPFC reduces task-related neuronal firing (Wang et al., 2007). This finding raises the question of how activation of HCN channels, which normally produces depolarization and increased firing, reduces the firing of PFC pyramidal cells.

It is well established that the influx of cations such as Ca^2+^ or Na^+^ can lead to hyperpolarization or reduced excitability by activation of K^+^ conductances (Kaczmarek et al., 2017). Experiments with olfactory mitral cells have found that Na^+^ entry through HCN channels stimulates K^+^ channels activated by intracellular Na^+^, termed K_Na_ channels (Lu et al., 2010). The two major known K_Na_ channels have been termed Slack and Slick (also K_Na_1.1 and K_Na_1.2, encoded by the *KCNT1* and *KCNT2* genes respectively. (Joiner et al., 1998; Bhattacharjee et al., 2003; Yuan et al., 2003; Kaczmarek et al., 2017). Although both are very widely expressed in the nervous system, the only cortical region in which immunoreactivity for Slack-B, a major splice isoform, is detected is the frontal cortex (Bhattacharjee et al., 2002). Human mutations that alter the function of Slack channels result in several distinct childhood epilepsies, all of which are associated with very severe intellectual impairment (Kim and Kaczmarek, 2014). Moreover, in one of these conditions (ADNLFE, autosomal dominant nocturnal frontal lobe epilepsy) seizures are restricted to the frontal lobes (Heron et al., 2012; Kim et al., 2014; Moller et al., 2015).

We have tested the hypothesis that Na^+^ influx through HCN channels activates Slack-B channels. We have found that Slack channels co-immunoprecipitate with HCN1, and that HCN1 and Slack channels colocalize on the same dendritic spine of mPFC pyramidal cells. ZD7288, a specific blocker of HCN channels, reduces K_Na_ current in pyramidal cells but has no effect on Slack channels when they expressed without HCN1. Co-expression of Slack-B with HCN channels reverses the effects of HCN activation on cytoplasmic Ca^2+^ levels. Finally, we describe a novel Slack channel blocker and demonstrate that it mimics and occludes the effect of HNC channel block on working memory performance.

## Results

### Slack channels complex with HCN channels in mouse mPFC neurons

To determine if Slack channels interact with HCN channels, we first carried out co-immunoprecipitation (Co-IP) experiments using extracts of mouse cerebral cortex. Immunoprecipitation was carried out using a previously described antibody against the cytoplasmic N-terminal domain that is specific to the Slack-B isoform that was generated (Bhattacharjee et al., 2002; Brown et al., 2008). An HCN1 channel band was readily detected in the Slack-B immunoprecipitates, but not in those carried out using control non-immune IgY antibodies (Fig. 1A).

**Figure 1.**
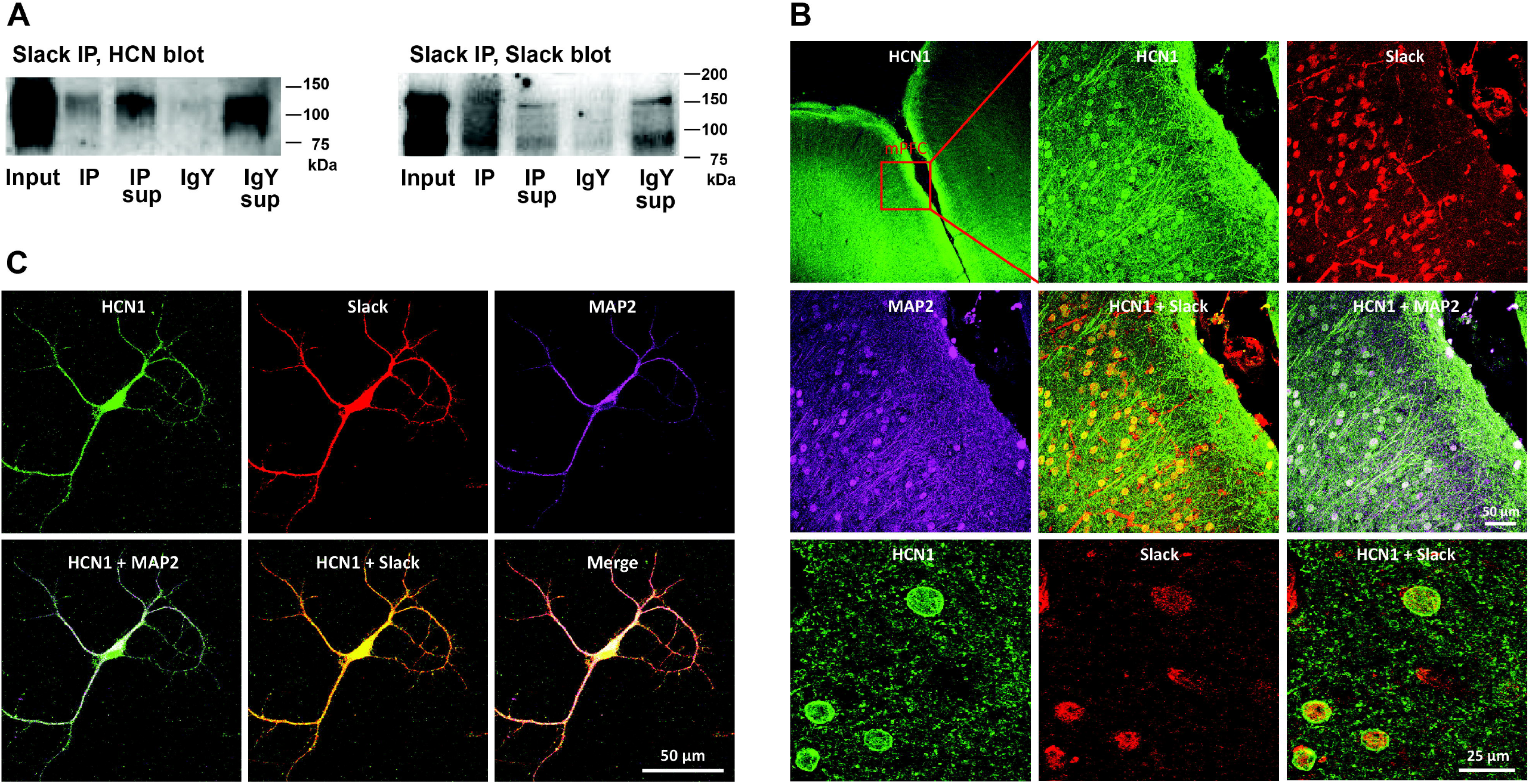
Slack channels co-immunoprecipitate and colocalize with HCN channels in mouse frontal cortex. **A**. Co-immunoprecipitation of Slack and HCN channels from mouse frontal cortex. Frontal cortex lysates were subjected to immunoprecipitation using anti-Slack-B antibody or chicken IgY, followed by Western blotting with either Anti-HCN1 or Anti-Slack antibody. Input: sample before Slack or IgY immunoprecipitation; IP: Slack IP sample; IP Sup: supernatant after Slack IP; IgY: IgY IP Sample; IgY sup: supernatant after IgY IP. *Left:* The presence of HCN1 band in the Slack IP sample, but not in the IgY IP lane indicates interaction of HCN1 channel with Slack channel. The presence of HCN1 in the IgY sup confirms no binding of the negative IgY beads with HCN1. *Right:* The presence of Slack band in the IP lane but absence in the IgY lane, indicating specificity of Slack antibody immunoprecipitation. **B**. Immunolocalization of Slack and HCN1 in 2-month-old mouse mPFC. Triple immunolabeling experiments depicting colocalization between HCN1 (green) and Slack (red) at the somata (MAP2, purple) of cells in layer II-III of mouse mPFC. Scale bar = 50 μm for all the photographs except for the first one and the last three (scale bar = 25 μm). **C**. Immunolocalization of Slack and HCN1 in cultured frontal cortical neurons on DIV14. Triple immunolabeling experiments depicting colocalization between HCN1 (green) and Slack (red) at the soma and dendrites (MAP2, purple) of cultured cortical neuron. Scale bar, 50 μm.

To further characterize the interaction between Slack and HCN channels, we carried out immunolocalization of these channels in mPFC (Fig. 1B) of 2-month-old mice and in primary cortical cultures on DIV 14 obtained from the frontal cortex of E16-17 mouse embryos (Fig. 1C). As described previously, we found that HCN1 immunoreactivity colocalized with KCNT1 in both mPFC neurons of mouse brain (Fig. 1B) and cultured cortical neurons (Fig. 1C). Notably, the colocalization of HCN1 and Slack channels was observed at the dendritic spine membranes of cultured neurons (Fig. 1C), which suggests that Slack channels may complex with HCN1 channels at dendritic spines of mouse mPFC neurons.

The immunolabeling experiments were also performed in Slack^−/-^/Slick^−/-^ mice. Robust expression of Slack-B channels was detected in pyramidal neurons in layers II-III of wild-type mouse mPFC (Fig. 1S). However, no immunolabeling was detected in cortices from Slack^−/-^ /Slick^−/-^ mice (Fig. 1S).

### Slack and HCN1 channels are co-localized within dendritic spines in rat mPFC layer II/III

We further conducted post-embedding immunoEM to determine the subcellular localization of HCN1 and Slack (Slo2.2, KCNT1) channels in rat prelimbic mPFC layer II/III, the cortical sublayers most associated with working memory microcircuits in primates. We utilized high-resolution, non-diffusible gold immunoprobes to precisely localize HCN1 and Slack channels on dendritic spine membranes, and determine their position in relationship to synaptic specializations. Immunoreactivity for HCN1 in rat mPFC layer II/III was observed in perisynaptic compartments near axospinous glutamatergic-like asymmetric synapses (Fig. 2A). An identical configuration has been previously observed in primate dorsolateral PFC layer III circuits, where HCN1 channels are visualized on the perisynaptic annulus of asymmetric synapses within dendritic spines, embedded at the edge of the postsynaptic density (PSD) (Paspalas et al., 2013a). Similarly, immunoreactivity for Slack channels was observed along the plasmalemma within dendritic spines, immediately next to asymmetric, presumed excitatory synapses in rat mPFC layer II/III (Fig.2B). In order to determine the spatial interactions of these channels and whether they are localized within the same dendritic spine subcompartment, we conducted dual-labeling immunogold immunoEM. Both HCN1 and Slack channels were co-localized at dendritic spine head membranes, perisynaptically next to glutamatergic synapses (Fig. 2C). These findings suggest that HCN1 and Slack potassium channels are strategically positioned on dendritic spines to gate excitatory synapses in mPFC layer II/III circuits.

**Figure 2.**
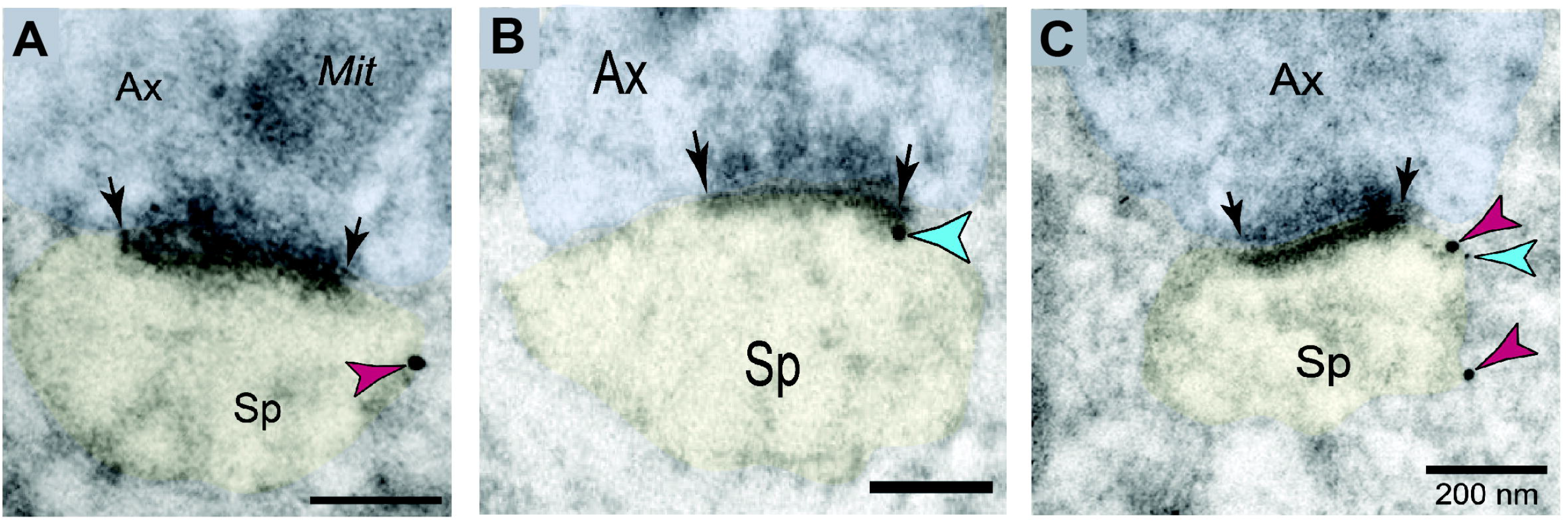
Co-localization of HCN1 and Slack channels within dendritic spines in rat mPFC layer II/III. **A**. Immunogold labeling (15 nm particles) reveals precise labeling of HCN1 within dendritic spines of pyramidal cells in rat prelimbic mPFC layer II/III. HCN1 channels are concentrated in the perisynaptic compartment adjacent to asymmetric, presumed glutamatergic-like synapses. **B**. Similar to HCN1 channel labeling, immunogold labeling (15 nm particle) for Slack channels is enriched along the plasma membranes bordering asymmetric axospinous synapses, receiving presumed glutamatergic input, within dendritic spines in rat mPFC layer II/III. **C**. Co-localization of HCN1 (15 nm particle) and Slack (5 nm particle) channels within dendritic spines in rat mPFC layer II/III. In dendritic spine heads, the immunoparticles for HCN1 and Slack overlap at perisynaptic locations. Synapses are between arrows. Color-coded arrowheads point to HCN1 (red) and Slack (blue) immunoreactivity. Profiles are pseudocolored for clarity. Ax, axon; Sp, dendritic spine; Mit, mitochondria. Scale bars, 200 nm.

### Coexpression of Slack with HCN channels in HEK cells reverses HCN-induced depolarization

To test whether the coupling of Slack-B to HCN channels reverses the normal physiological effects of HCN channel activation, we next carried out experiments using a previously described, synthetic excitable cell type (Thomas and Hughes, 2020). These HEK cells, termed Kuhl-H cells, express HCN channels, which are activated by increases in cAMP levels, as well as a light-sensitive adenylyl cyclase enzyme (bPAC). Thus, exposure of the cells to blue light triggers opening of the HCN channels, resulting in a depolarization of the cells that produces an increase in cytoplasmic Ca^2+^ levels. The cells additionally express a Ca^2+^ sensor (R-GECO1), allowing monitoring of the effects of HCN channel activation in live cells in real time. The functional components within these cells are illustrated in Figure 3A.

**Figure 3.**
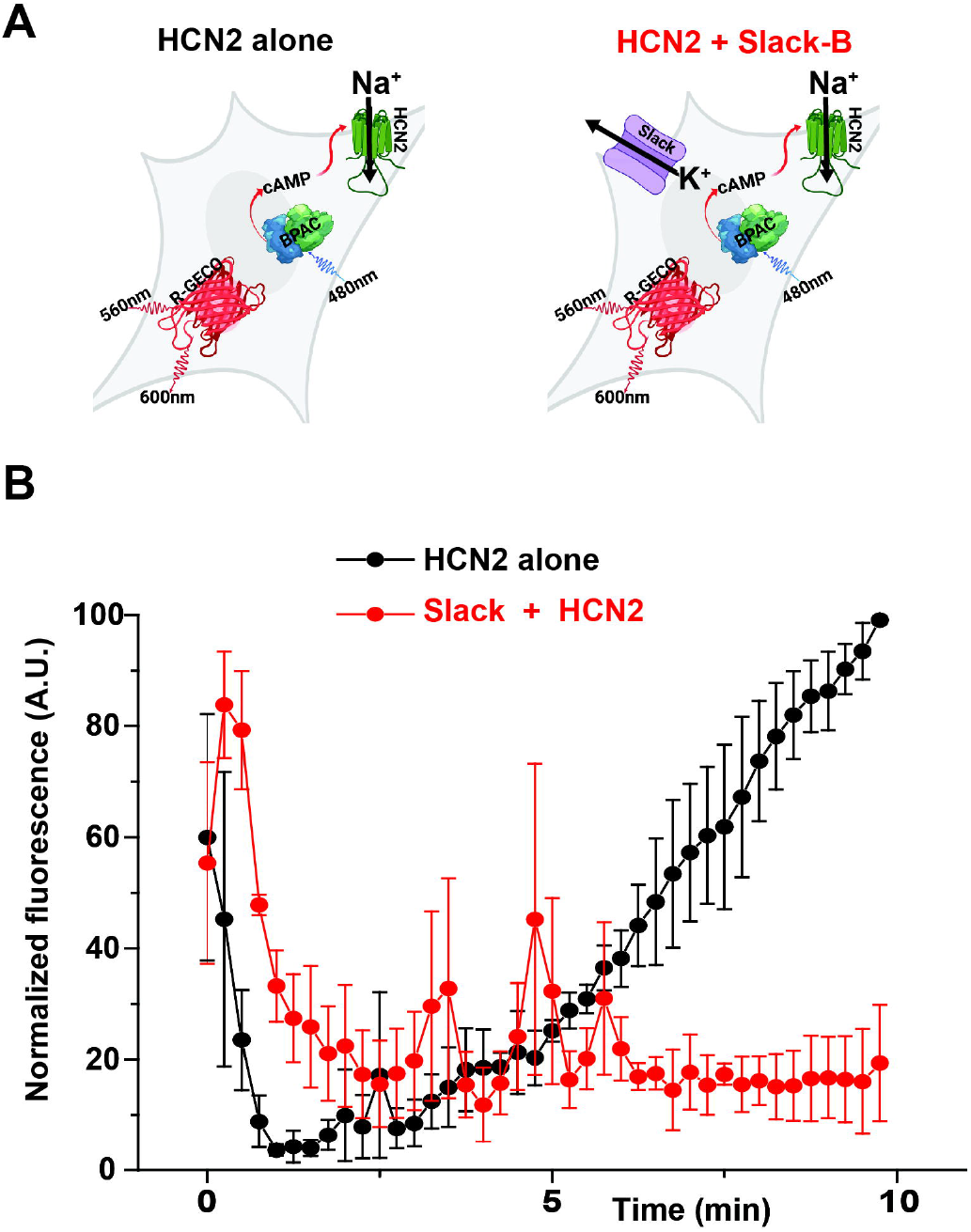
Slack-B channels reverse the cellular effects of HCN channel activation. **A**. Diagram of components in HEK cells used for the assay. HEK-293 cells are transduced with bPAC (the actuator), HCN2 (ion channel that is cAMP coupled), and R-GECO1 (a Ca^2+^ sensor). Brief pulses of blue light (50 ms of 488nm light) activate bPAC to increase cAMP levels that results in opening of the HCN2 channel, producing a slow increase in Ca^2+^ fluorescence signal (Thomas and Hughes 2020). **B**. Responses to light pulses in cells expressing HCN2 alone with those co-expressing Slack-B with HCN2 channels. Combined plots of normalized data comparing the responses of cells expressing HCN2 alone with those co-expressing Slack-B with HCN2 channels. Results are show mean ± SEM for three independent experiments each of which measured responses of 52 independent cells for a total of 156 cells in each condition.

As described previously, in the absence of Slack channels, application of brief pulses of blue light to these cells elevates cAMP and activates the HCN channels to depolarize the cells, resulting in a slow increase in Ca^2+^ reporter fluorescence (Thomas and Hughes, 2020) (Fig. 3B). Elavations of cAMP have been found to have no effect on Slack-B channels expressed in heterologous systems (Nuwer et al., 2009). When, however, Slack-B was coexpressed with the HCN channels in the Kuhl-H cells, the response to the pulses of blue light was substantially reduced or abolished (Fig. 3B). These findings are consistent with the hypothesis that the coupling of HCN channels to Slack-B reverses the depolarization normally produced by HCN activation.

### Inhibition of HCN channels reduces K_Na_ currents in mPFC Pyramidal Neurons

To test further the hypothesis that HCN channel activation leads to the activation of K_Na_ potassium channels, we carried out whole cell voltage clamp recordings on primary cortical neurons and on mPFC pyramidal neurons in cortical slices, and tested the effects of the HCN channel blocker ZD7288 on K_Na_ currents induced by voltage steps. We found that treatment with ZD7288 10 μM produced a marked reduction in outward K^+^ current in primary cortical neurons (Fig. 4A-4C) as well as in mPFC pyramidal neurons of intact mice (Fig. 4D-4F). However, the ZD7288 10 μM did not produce any reduction in K^+^ current in the primary cortical neurons when blocking Slack channels with a novel Slack blocker, SLK-01 (Fig. 4G and 4H, see next section).

**Figure 4.**
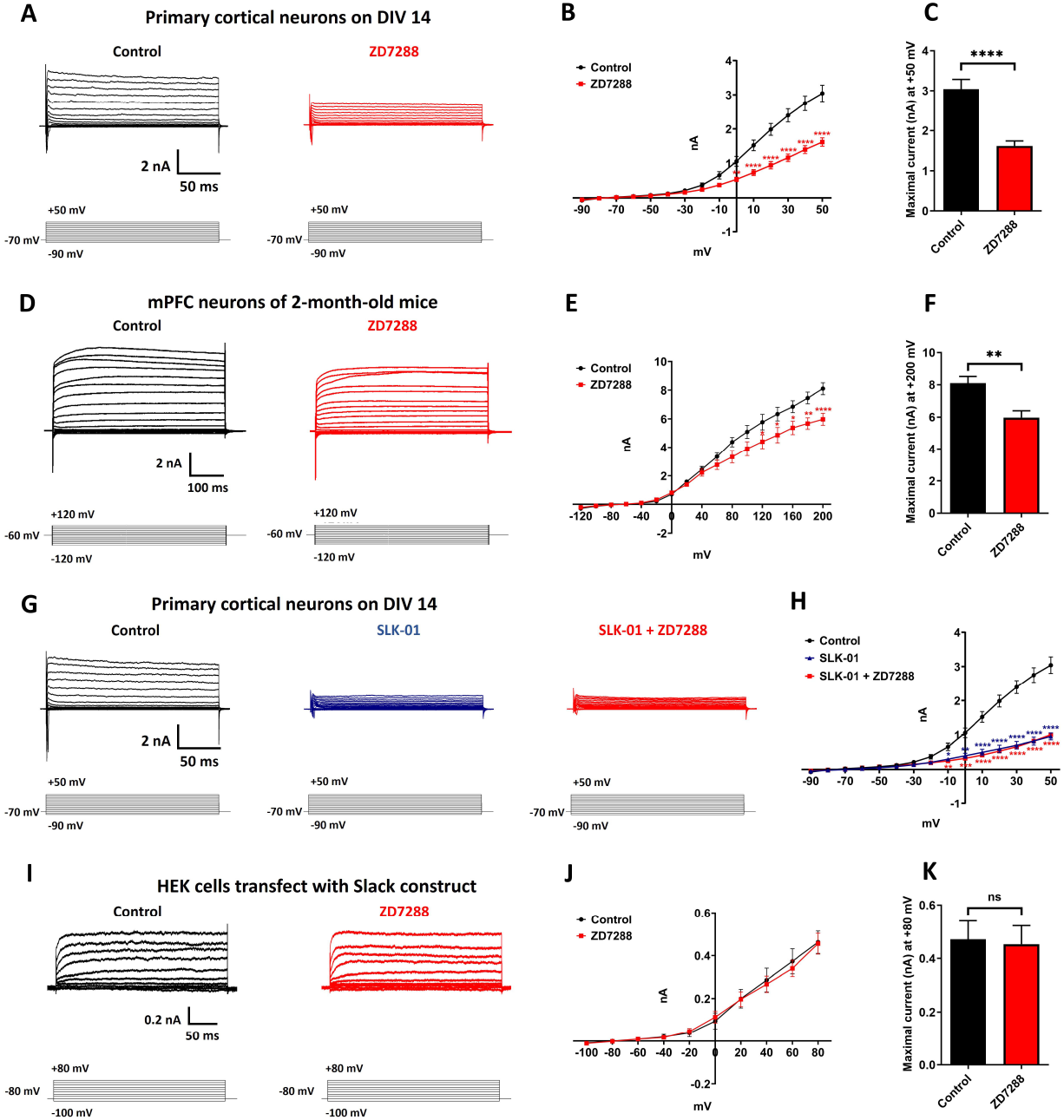
Blocking HCN channels with ZD7288 reduces outward K^+^ currents in mPFC pyramidal neurons. **A**. Representative current traces from whole cell voltage clamp recordings in cultured frontal cortical neurons on DIV14 induced by voltage steps from -90 to +50 mV in 10 mV increments before and after application of HCN channel blocker ZD7288 (10 μM, 10 min). **B**. Summary data show the outward K^+^ currents for each voltage step in neurons before and after application of ZD7288. Data are shown as mean ± SEM (n = 12-14, two-way ANOVA). **C**. Maximal K^+^ current at +50 mV for each condition. Data are shown as mean ± SEM (n = 12-14, Student’s t test). **D and E**. Representative current traces and summary data from whole cell voltage clamp recordings in mPFC pyramidal neurons of 2-month-old mice induced by voltage steps from -120 to +200 mV in 20 mV increments before and after application of ZD7288 (10 μM, 10 min). Data are shown as mean ± SEM (n = 6, two-way ANOVA). **F**. Maximal K^+^ current at +200 mV for each condition. Data are shown as mean ± SEM (n = 6, Student’s t test). **G and H**. Representative current traces and summary data from whole cell voltage clamp recordings in cultured frontal cortical neurons on DIV14 induced by voltage steps from -90 to +50 mV in 10 mV increments before and after application of Slack channel blocker SLK-01 (10 μM, 10 min) or SLK-01 with ZD7288 (10 μM, 10 min). Data are shown as mean ± SEM (n =7-12, two-way ANOVA). **I and J**. Representative current traces and summary data from whole cell voltage clamp recordings in HEK cells stably expressing Slack induced by voltage steps from -120 to +80 mV in 20 mV increments before and after application of ZD7288 (10 μM, 10 min). Data are shown as mean ± SEM (n = 4, two-way ANOVA). **K**. Maximal K^+^ current at +80 mV for each condition. Data are shown as mean ± SEM (n = 4, Student’s t test). \**p <* 0.05, \*\**p <* 0.01, \*\*\**p* *<* 0.001 and \*\*\*\**p <* 0.0001.

The suppression of outward current by ZD7288 in neurons could in theory results from an off-target effect of this compound on K_Na_ channels. To test this hypothesis, we carried out whole cell voltage recordings on HEK cells stably expressing the Slack channels. We found that addition of the HCN inhibitor, ZD7288 (10 μM) to the external medium had no effect on Slack currents in this cell line (Fig. 4I-4K). Collectively, these results suggest that the reduction in K^+^ current in mPFC pyramidal neurons by ZD7288 is mediated by its inhibition of HCN channels.

### SLK-01 is a use-dependent blocker of Slack Channels

Known pharmacological agents that have been used previously to inhibit Slack channels, such as quinidine (Yang et al., 2006), have significant non-specific effects on additional ion channels, including other K-channels. In an attempt to further probe the interaction of HCN channels with Slack in intact animals, we have now characterized a novel use-dependent Slack blocker. The structure of this compound, SLK-01 (originally given the identifier BMS-271723, see Discussion) is shown in Figure 5A. In the present study, currents were recorded from HEK cells stably expressing the Slack channel using the whole cell patch clamp technique. SLK-01 was applied to these cells at concentrations of 50 nM to 40 μM (n = 7, Fig. 5B and 5C). At concentrations above 1 μM there was a concentration-dependent reduction in current amplitude accompanied by a change in kinetic behavior. On depolarization of untreated cells, Slack currents activate monotonically to a final current level that persists throughout the duration of the depolarizing command. In contrast, in the SLK-01 treated cells, the currents in response to command potentials greater than +20 mV rose to a peak within 50 msec followed by a rapid decline to a lower level (Fig. 5B). The amplitude of the peak current in the presence of SLK-01 to the maximal current in untreated cells was reduced with an IC_50_ of ∼20 μM (Fig. 5C). The voltage dependence of Slack currents was, however, not altered by SLK-01(Fig. 5D). The carrier solution had no effect on Slack currents.

**Figure 5.**
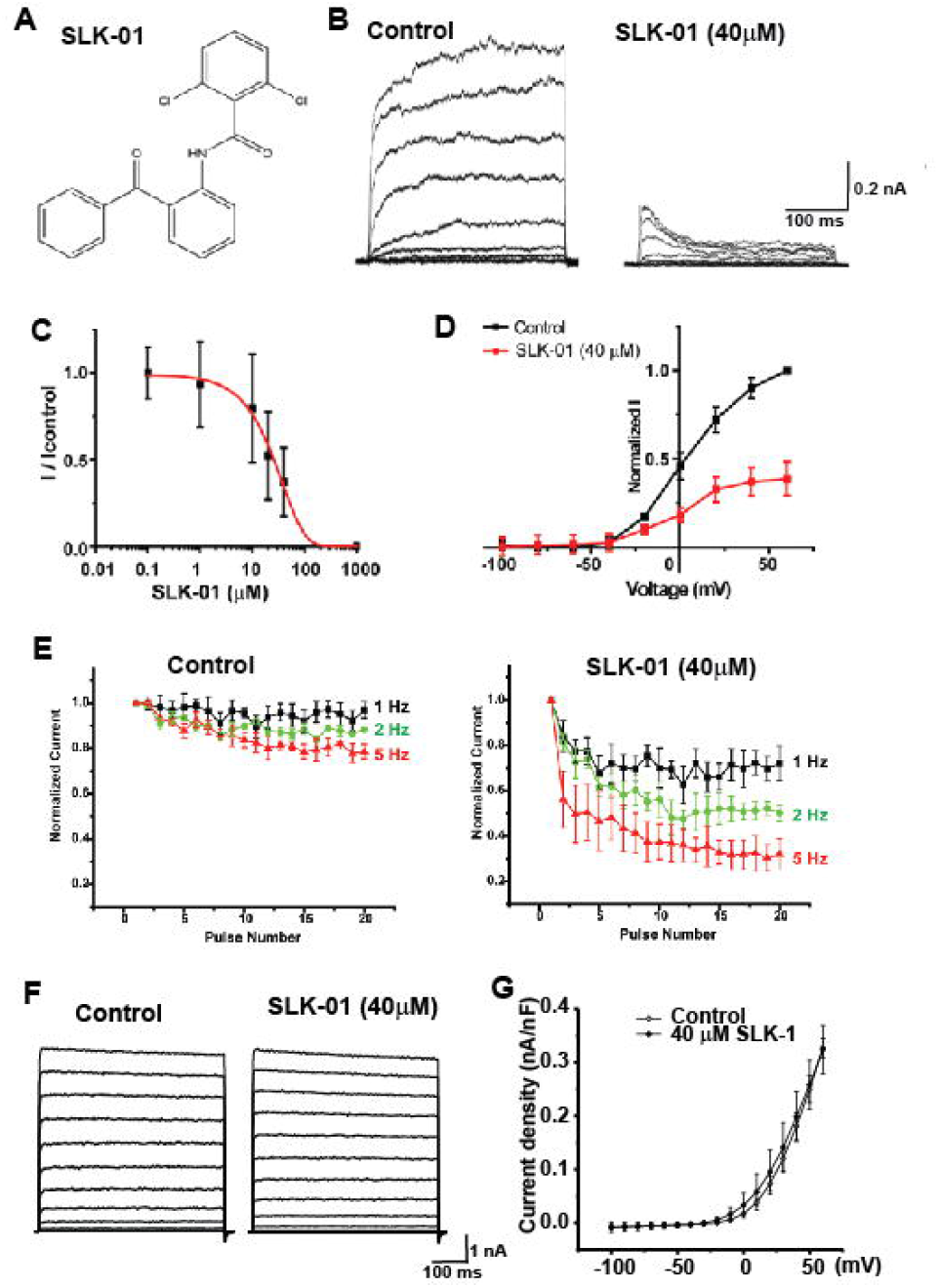
SLK-01 inhibits Slack currents. **A**. Structure of SLK-01. **B**. Representative traces of currents elicited by voltage steps between -100 and +60 mV from a holding potential of -80 mV, in control medium and afer application of 40 μM SLK-01. **C**. Concentration-response relationship for SLK-01 at +40 mV. Compound was sequentially applied to cells (n = 7) and the peak current measured. The inhibition increased during depolarizing commands producing near complete blockade after 50 ms at the highest concentration (40 μM). **D**. Current-voltage relationships for cells treated with 40 μM SLK-01. **E**. Use-dependence of block of Slack currents by SLK-01. Currents were evoked by step depolarizations (220 ms) from a holding potential of - 80 mV to +40 mV at rates of 1, 2 or 5 Hz. Plots show the progressive change in current amplitude recorded at the end of each pulse in the absence (n = 4) or presence of 40 μM SLK-01 (n = 3). **F**. SLK-01 has no effect on BK currents. Representative traces of currents in CHO cells stably expressing rat BK channels before and after application of 40 μM SLK-01. Currents were evoked by voltage steps between -100 and +60 mV from a holding potential of -80 mV. **G**. Group data quantifiying the mean amplitude of currents recorded at +60 mV before and after SLK-01 (20 μM, n = 4; 40 μM n = 6).

The decline in Slack currents following their activation in the presence of SLK-01 suggest that block of the channels by this agent is use dependent, such that block only occurs after channels have been opened in response to depolarization. If this is the case, repeated depolarization at a high rate would be expected to promote the rate at which channels become blocked. To test this, we applied trains of 20 or 100 depolarizing commands from holding potential of -80 mV to 40 mV for 220 ms at different rates and measured the degree of block at the end of each pulse. The percent inhibition for each pulse was normalized to that for the first pulse in each series. In untreated cells, repeated stimulation at 1or 2 Hz produced no change in current amplitude throughout the train, and stimulation at 5 Hz produced an only a small reduction in current amplitude after 20 pulses (∼20%, Fig. 5E). After treatment with 40 μM SLK-01, the amplitude of Slack current measured at the end of each pulse progressively decreased throughout the train even at 1 Hz (Fig. 5E), and at 5 Hz, currents were suppressed by ∼70%, confirming the use-dependence of block by this agent.

As a partial test for specificity, we measured the effects of SLK-01 on BK potassium currents. BK large-conductance calcium-activated channels (KCNMA1, K_Ca_1.1, Slo1) are the channels most closely related to Slack, other than its paralog Slick (KCNT2, K_Na_1.2) (Joiner et al., 1998; Bhattacharjee et al., 2003). Application of either 20 μM or 40 μM SLK-01 to cells stably expressing BK channels had no statistically significant effect on the amplitude of BK currents (Fig. 5F, G).

### SLK-01 infusion into PFC improves delayed alternation performance, similar to HCN channel blockade

Previous work has demonstrated that infusion of a very low dose of the HCN channel blocker, ZD7288, into rat PFC improved delayed alternation performance of rats (Wang et al., 2007), the classic test for assessing working memory in rodents that is dependent on the integrity of the mPFC. These behavioral data were consistent with physiological data from monkeys, where low, but not high, doses of ZD7288 enhanced delay-related neuronal firing (Wang et al., 2007). In the delayed alternation task in a T maze, the rat must remember which arm it had chosen on the previous trial, and alternate its response to the other side to receive reward (Fig. 6A). This requires both spatial working memory and response inhibition to overcome the tendency to go to a previously rewarded location, both functions of the PFC. If a major mode of action of HCN channels on mPFC pyramidal neurons is to activate Slack channels, the inhibition of Slack channels would be expected to have a similar effect to that of HCN blockade. To test this possibility, rats (n=5) were implanted with indwelling cannula above the prelimbic PFC, and were infused with vehicle or SLK-01 (20 μM or 50 μM in 0.5μl/side). While there were no effects seen with the 20μM dose, 3 of the 5 rats were improved by the 50μM dose, and 2 were impaired. As blockade of ion channels often has an inverted U dose-response, we tested an intermediate dose, 30μM, in the two rats that were impaired by the 50μM dose, and found that they were improved compared to vehicle control (Fig. 6B). Thus, a dose of either 30 μM or 50μM SLK-01 was able to significantly improve performance compared to vehicle control (best dose vs. vehicle: p<0.009). As Slack channel blockade mimics the effects of HCN channel blockade, these behavioral data are consistent with the hypothesis that the activation of HCN channels is coupled to Slack channel opening.

**Figure 6.**
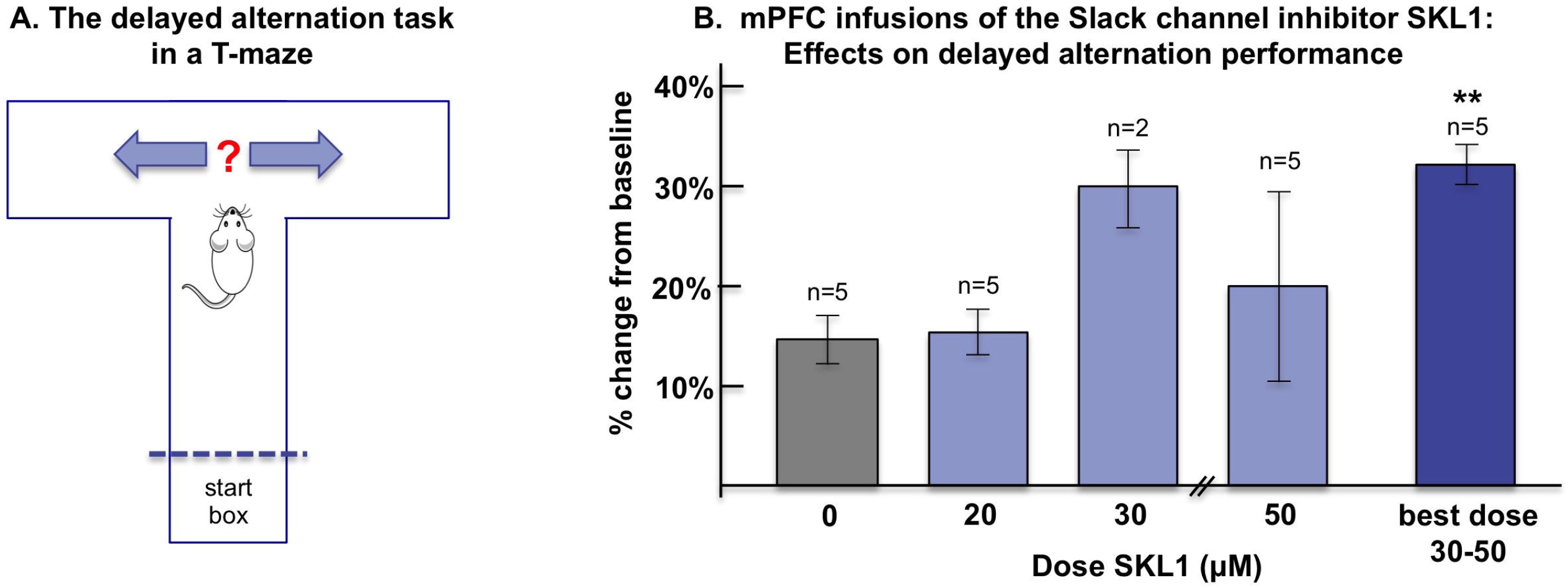
Blockade of Slack channels with infusion of SLK-01 into rat mPFC improved delayed alternation performance in a T maze. **A**. Diagram of the delayed alternation task in a T maze for testing spatial working memory performance in the rat. On the first trial, the rat can choose either arm and be rewarded, on subsequent trials the rat must alternate its response. The rat spends the delay period in the start box, and thus must remember its previous spatial response over this delay. **B**. Infusion of SLK-01 (20-50 μM) produced a dose-related improvement in working memory performance, where a best dose of either 30 or 50 μM significantly improved performance compared to vehicle control (\*\**p* < 0.009, one-way ANOVA).

## Discussion

We have found that Slack-B channels form a complex with HCN channels in pyramidal cells of the rodents mPFC. A clear implication of this finding is that Na^+^ entry through HCN channels serves to activate the Na^+^-dependent K^+^ current of the Slack channel. Because the reversal potential of the non-selective HCN cation channels is around -20 mV, activation of HCN channels alone would be expected to depolarize neurons under physiological conditions, leading to increased excitability. Coupling of HCN to Slack K_Na_ currents, however, would be expected to suppress the depolarization and result in a hyperpolarization and/or a decrease in input resistance leading to decreased membrane excitability. Consistent with this hypothesis, we found that blockade of HCN channels with ZD7288 reduces K^+^ current in pyramidal cells, but has no effect on K^+^ currents in a cell line stably expressing Slack alone. Finally, infusion of the Slack channel blocker, SLK-01, into rat mPFC improved working memory performance, occluding the beneficial effects of the HCN channel blocker, suggesting the two agents may act through a common pathway.

Our findings may resolve the paradox that activation of HCN channels in PFC leads to a loss of task-related firing (Wang et al., 2007) while low dose blockade of HCN channels enhances task-related firing (Wang et al., 2007; Wang et al., 2011). The ability to carry out spatial working memory tasks requires persistent firing of pyramidal cells arising from the recurrent excitation of layer III microcircuits that make synapses on spines (Goldman-Rakic, 1995). These layer III spines express HCN channels (Wang et al., 2007; Paspalas et al., 2013b), and as shown in the current study, they can be co-expressed with Slack channels on dendritic spines in rat mPFC. Thus, increased cAMP opening of HCN channels in spines would lead to Slack channel opening, spine hyperpolarization, and the loss of recurrent excitation needed to sustain working-memory related neuronal firing. This is in contrast to the more classic HCN channel actions on distal dendrites where they are distant from synapses, and depolarize the membrane when the neuron becomes hyperpolarized, thus having an important excitatory influence (He et al., 2014). Layer V cortical pyramidal cells often express high levels of HCN channels on their distal dendrites, including in rodent mPFC (Lörincz et al., 2002; Notomi and Shigemoto, 2004; Day et al., 2005), indicating circuit selective actions of these channels dependent on their location and molecular interactions.

A required role of Slack channels in the control of cortical functions such as working memory is expected based on the known effects of Slack mutations. Within the cerebral cortex, Slack-B, a major splice isoform is expressed selectively in the frontal cortex (Bhattacharjee et al., 2002). Human mutations that alter the function of Slack channels result in several distinct childhood epilepsies, all of which are associated with very severe intellectual impairment (Kim and Kaczmarek, 2014). When expressed in heterologous systems, the majority of disease-causing mutations give rise to Slack currents that are increased in amplitude from 3-22-fold over those of wild type channels, with no change in levels of protein or RNA stability (Kim et al., 2014; Martin et al., 2014). One of the disease caused by Slack mutations is EIMFS (Epilepsy of Infancy with Migrating Focal Seizures) (Barcia et al., 2012; Ishii et al., 2013; McTague et al., 2013; Shimada et al., 2014; Moller et al., 2015; Ohba et al., 2015; Rizzo et al., 2016) and over 50% of patients with MMPSI have been documented to have Slack mutations (Barcia et al., 2012; Kim and Kaczmarek, 2014; Fleming et al., 2016). Seizures in MMPSI typically begin within the first few weeks after birth and diminish with maturity. A second condition resulting from Slack mutations is ADNLFE (autosomal dominant nocturnal frontal lobe epilepsy) (Heron et al., 2012; Kim et al., 2014; Moller et al., 2015). In this condition, seizures begin at 6-8 years of age, are restricted to the frontal lobes and occur only at night. Slack mutations are also found in other seizure disorders (Juang et al., 2014; Vanderver et al., 2014; Moller et al., 2015; Ohba et al., 2015; Hansen et al., 2017), including Ohtahara syndrome(Martin et al., 2014). Finally, as many as 40% of patients with severe autism have early-onset seizures that disappear later in development (Musumeci et al., 1999; Muhle et al., 2004; Amiet et al., 2008; Bailey et al., 2008; Yasuhara, 2010; Jokiranta et al., 2014). Slack mutations have also been documented for autism (Iossifov et al., 2014).

As part of this study, we characterized a novel compound, SLK-01, that is an effective use-dependent inhibitor of Slack channels. This compound was originally synthesized as part of a program targeting KCNQ2/3 and BK (*Slo1)* channels at the Bristol-Myers Squibb (BMS) Pharmaceutical Research Institute in the Department of CNS Drug Discovery in the late 1990s/early 2000s and given the identifier BMS-271723. It was initially screened against both the primary target and select secondary K channels. During the course of this screening it was discovered that some of these lead compounds did not significantly activate KCNQ or BK channels, but possessed inhibitory activity at Slack channels. The modulatory action of ∼10 of these compounds was determined using heterologous expression of Slack in *Xenopus* oocytes and was presented in a discussion of off-target activity discovered in compounds originally synthesized as KCNQ2/3 and BK channel modulators (Gribkoff et al., 2004).

Our findings indicate that SLK-01 is an open-channel blocker, suggesting that it could be more potent on human disease-relevant mutant gain-of-function Slack channels than on wild-type channels (Barcia et al., 2012; Heron et al., 2012; Kim and Kaczmarek, 2014; Milligan et al., 2014; Moller et al., 2015). Until very recently, the only other effective compounds known to block Slack channels was quinidine, an agent that also acts on a wide variety of other channels (Yang et al., 2006). In the past year, there have been reports of other more selective agents that block Slack channels (Cole et al., 2020; Spitznagel et al., 2020; Griffin et al., 2021; Qunies et al., 2022). Further studies will be required to determine the specificity, efficacy and species selectivity of these compounds, as well as that of SLK-01.

In summary, our studies strongly suggest that a complex containing both Slack and HNC channel plays a key regulatory role in the intrinsic excitability of medial prefrontal pyramidal cells of the cerebral cortex and in the response of their dendrites to incoming stimuli. that are required. Further genetic studies will be required to establish the specific function of this two-channel complex in the regulation of working memory performance.

## Acknowledgements

This work was supported by NIH grants NS102239 (LKK), DC01919 (LKK), P1AG047744 (AFTA), 1RL1AA017536 (AFTA), U54RR024350 (AFTA) and by FRM DEQ20170336753. We thank Mark Plummer and Denton Hoyer of the Yale Center for Molecular Discovery for the synthesis of SLK-01.

## Materials and methods

All experiments were conducted in accordance with the guidelines of Yale University Institutional Animal Care and Use Committee, and Public Health Service requirements for animal use as described in the Guide for the Care and Use of Laboratory Animals.

### Primary Cortical Neuron Culture

Primary cortical neurons were prepared from E16-17 mouse embryos as described previously with modifications specific for this study (Park et al., 2017). After isolation of frontal cortex from embryonic brains, neurons were dissociated and seeded (on coverslips inside a 6 well plate: 0.2E6 cells/well) onto plates containing NB plus [Neurobasal medium supplemented with B27 (Invitrogen GIBCO Life Technologies), GlutaMAX (GIBCO), and penicillin/streptomycin (GIBCO)] and 5% FBS (GIBCO). After 2 hours incubation, primary cultures were maintained in NB plus without FBS in a 5% CO_2_ and 20% O_2_ incubator at 37 °C. Subsequently, half the medium was replaced every 2 days.

### Antibodies

HCN1 channel immunolabeling used three antibodies: (1) mouse monoclonal antibody to HCN1 obtained from NeuroMab/Antibodies Incorporated (catalog# 75-110), (2) mouse monoclonal antibody to HCN1 obtained from Millipore-Sigma (catalog# SAB5200024, clone S70-28), and (3) rabbit polyclonal antibody HCN1 obtained from Abcam (catalog# ab229340). Slack (Slo2.2, KCNT1) channel immunolabeling used a previously generated affinity-purified custom polyclonal chicken antibody against the N-terminal of rat Slack (Bhattacharjee et al., 2002) and a previously validated mouse anti-Slack cytoplasmic, C-terminal antibody (Shore et al., 2020; Gertler et al., 2022).

### Tissue preparation

For HCN1 and Slack channel immunolabeling, 3 male, 27 to 29-month-old Sprague-Dawley rats and 3 male 2-month-old C57BL/6J mice were used. Aged rats were used due to potentially increased HCN1 actions mediated by disinhibited cAMP-PKA signaling in the mPFC with advancing age (Ramos et al., 2003). Animals were anesthetized with Nembutal (50mg/mL, i.p.) and perfused transcardially with 4% paraformaldehyde (PFA), 0.05% glutaraldehyde in 0.1M phosphate buffer (PB; pH7.4). After perfusion, brains were removed from the skull and immersed in 4% PFA overnight at 4 °C. Coronal 60 μm-thick sections were then cut on a Vibratome (Leica V1000) and collected in 0.1 M PB.

### Immunofluorescence

For immunocytochemistry of primary cortical neurons on DIV 14, coverslips were fixed in 4% paraformaldehyde for 10 min and permeabilized in blocking buffer (0.2% Triton X-100 and 3% normal goat serum in PBS) for 5 min at room temperature. For immunohistochemistry of cerebral cortex from 2-month-old mice, brain slices were permeabilized and incubated in blocking buffer for one hour at room temperature. After blocking, sections from DIV 14 cortical neuronal cultures or 2-month-old mice were overlaid with primary antibodies to Slack (1:200, Neuromab), HCN1 (1:200, Thermo Fisher Scientific) and MAP2 (1:200, Santa Cruz Biotechnology) overnight at 4 °C. Then, the corresponding Alexa Fluor 488-, 546- or 647-conjugated secondary antibodies were applied. Stained sections were mounted with DAPI-containing mounting solution and sealed with glass coverslips. Confocal imaging of primary cortical neurons and brain slices was carried out on a Leica SP5 MP. Three-dimensional z-stack images were taken and stitched into tiles to cover the entirety of the neurons of interest.

### Post-embedding immunogold immunohistochemistry

In order to visualize the subcellular localization and interaction between HCN1 and Slack (Slo2.2, KCNT1) channels in rat mPFC layer II/III circuits, we utilized post-embedding immunogold dual-labeling techniques. Briefly, frozen brain tissue was freeze substituted in a Leica EM AFS2 unit starting from -90°C, and gradually increasing temperature to -45°C. After 3 changes of acetone, tissue was infiltrated with Lowicryl HM20 resin. The polymerization was carried out at -20°C with ultraviolet. Hardened blocks were cut using a Leica UltraCut UC7.

Sixty nanometer sections were collected on formvar/carbon-coated nickel grids and stained using 2% uranyl acetate and lead citrate. For immunolabeling of resin sections, grids were placed section side down on drops of 1% hydrogen peroxide for 5 min, rinsed, and blocked for nonspecific binding with 3% bovine serum albumin in Tris-buffered saline (TBS) containing 1% Triton X-100 for 30 min. For single immunogold labeling, grids were incubated with anti-HCN1 or anti-Slack primary antibody 1:100 overnight, rinsed in TBS then incubated with 15 nm protein A gold (UMC, Utrecht, The Netherlands) for 30 min. For gold-gold dual-labeling immunohistochemistry, grids were incubated with anti-HCN1 and anti-Slack primary antibodies 1:100 overnight, rinsed in TBS then incubated with 15 nm (HCN1) and 5 nm (Slack) protein A gold (UMC, Utrecht, The Netherlands) for 30 min. The grids were rinsed in PBS, fixed using 1% glutaraldehyde for 5 min, rinsed again, dried, and heavy metal stained using 2% aqueous uranyl acetate and lead citrate. Specimen preparation for electron microscopy was carried out by the Yale CCMI Electron Microscopy facility.

### Electron microscopy and data analysis

Tissue blocks containing prelimbic mPFC layer II/III were analyzed under a JEM1010 (Jeol, Tokyo, Japan) transmission electron microscope at 80 kV. Several blocks of each brain were examined and structures were digitally captured at x25,000-x100,000 magnification with Bioscan camera (Gatan, Pleasanton, CA, United States of America) and individual panels were adjusted for brightness and contrast using Adobe Photoshop and Illustrator CC.2017.01 image editing software (Adobe Systems Inc., San Jose, CA, United States of America). Approximately, 400 micrographs of selected areas of neuropil with immunopositive profiles were used for analyses. For profile identification, we adopted the criteria summarized in (Peters et al., 1991). *Western blot and Co-immunoprecipitation (Co-IP)*

Two-month-old mice frontal cortex were prepared using Pierce IP Lysis Buffer (Thermo Scientific) supplemented with cOmplete EDTA-free Protease Inhibitor Cocktail (Millipore Sigma) according to manufacturer’s protocol. Protein quantification was performed using Pierce BCA Protein Assay Kit (Thermo Scientific).

For Co-IP experiments, frontal cortex lysates were incubated with 5μg anti-Slack IgY (AvesLabs) antibody or IgY control (AvesLabs) overnight at 4°C. 100 μL Anti-IgY PrecipHen beads (AvesLabs) was added to sample and allowed to incubate for 2 hours, followed by wash and collection of beads. Beads were transferred to 2x Laemelli-Buffer with 5% beta-mercaptoethanol and incubated at room temperature for 30 minutes prior to Western blotting (Fleming et al., 2016).

For all immunoblotting experiments, protein samples were electrophoretically separated on an SDS-PAGE gel (4%–15% gradient gel, Bio-Rad) and transferred onto PVDF membranes (0.2 μm pores, Bio-Rad, USA). Blots were blocked in 5% nonfat milk in Tris-buffered saline and Tween 20 (TBST) for 1 h at room temperature and probed with the primary antibodies to Slack (1:3000, AvesLabs) and HCN1 (1:1000, Abcam) overnight at 4°C. After overnight incubation, the blots were washed three times in TBST for 30min, followed by incubation with corresponding horseradish peroxidase-conjugated secondary antibodies (1:1000; Abcam) at room temperature for 1 h. Protein bands were visualized via enhanced chemiluminescence and quantified with analyzed with ImageJ (NIH) software.

### Kuhl-H cells

HEK293 and HEK293 Slack-positive cells were cultured in DMEM, 10% FBS, and 1% Penicillin (100U/mL). For experiments, the cells were plated in 35mm plastic plates at 2ml per well (∼3×10^4^ cell/mL). To introduce components into the cells using baculovirus, HCN2 and the photoactivated cyclase, bPAC were packaged in BacMam (Montana Molecular, Bozeman, MT). The virus titers were bPAC (2 × 10^10^ VG/mL) and HCN2 (1.57 × 10^10^ VG/mL). The additional Ca^2+^ biosensor in baculovirus was obtained from Montana Molecular: R-GECO (2 × 10^10^ VG/mL). The cells were transduced following the manufacturer’s recommended protocol, plated in 35 mm plastic plates and transduced with the appropriate mix of viruses and HDAC inhibitor sodium butyrate (2mM final concentration). Two days later, the media was exchanged with PBS before imaging. Cells were imaged using post brief illumination from a blue LED (488nm) for rapid stimulation of bPAC. The yellow illumination was provided with a LED (560nm).

For data analysis, image data was stored in a Z-stack tiff file and loaded into the FIJI distribution of the IMAGEJ software. The cells were selected using a freehand ROI surrounding the cell of interest. The average total intensity pixel value within the ROI for each frame was collected using the time series analyzer plugin and saved as a .csv file. The raw fluorescence data was then put into PRISM and ORIGIN for further analysis. The fluorescence traces were rescaled from 0 - 100% and normalized to zero using PRISM software.

### Patch-clamp Recordings

Whole-cell patch-clamp recordings were performed with patch-clamp amplifiers (MultiClamp 700B; Molecular Devices) under the control of pClamp 11 software (Molecular Devices). Data were recorded with a sampling rate at 20 kHz and filtered at 6 kHz. Rs compensation of 70% was used. Primary cortical neurons at DIV 13-14 or mPFC pyramidal neurons in brain slices from 2-month-old mice or HEK cells were recorded at physiological temperature (37°C).

Recording electrodes were pulled from filamented borosilicate glass pipettes (Sutter Instrument, CA), and had tip resistances between 4 and 6 MΩ when filled with the following internal solution (in mM): 124 K-gluconate, 2 MgCl_2_, 13.2 NaCl, 1 EGTA, 10 HEPES, 4 Mg-ATP, and 0.3 Na-GTP (pH 7.3, 290-300 mOsm). The extracellular medium contained the following (in mM): either 140 NaCl, 5.4 KCl, 10 HEPES, 10 glucose, 1 MgCl_2_, and 1 CaCl_2_ (pH 7.4, 310 mOsm). For voltage-clamp recording on primary cortical neurons, neurons were held at -80 mV and given 60 ms voltage pulses in 10 mV steps over a range of -90 to +50 mV. For voltage-clamp recording on mPFC pyramidal neurons, neurons were held at -60 mV and given 500 ms voltage pulses in 20 mV steps over a range of -120 to +120 mV. For voltage-clamp recording on HEK cells, cells were held at -80 mV and given 300 ms voltage pulses in 20 mV steps over a range of -100 to +80 mV. ZD 7288 was purchased from MedChemExpress (Cat#: HY-101346) and was bath applied.

### SLK-1 Methods

SLK-01 was prepared and its structure confirmed by the Yale Center for Molecular Discovery. at Yale). Electrophysiological recordings were carried out on an HEK cell line stably expressing the human Slack channel. The cells were cultured in a modified low sodium DMEM medium supplemented with 10% fetal bovine serum and penicillin-streptomycin (Invitrogen Inc, Carlsbad, CA). Whole-cell patch recordings were obtained at room temperature (21-23 °C) using electrodes of 3-5 MΩ resistance. These were pulled from TW150F-6 micropipettes (World Precision Instruments Inc., Sarasota, FL) on a horizontal Flaming/Brown micropipette puller (Model P-87, Sutter Instrument Co., Novato, CA). During recordings, the cells were bathed in a solution containing (in mM): 140 NaCl, 1 CaCl_2_, 5 KCl, 29 glucose, and 25 HEPES, pH 7.4. The pipette solution contained (in mM): 100 K-gluconate, 30 KCl, 5 Na-gluconate, 29 glucose, 5 EGTA, 2 Na_2_ATP, 0.2 GTP, and 10 HEPES, pH 7.3. In general, currents elicited by voltage steps between -100 and +60 mV from a holding potential of -80 mV. SLK-01 stock solutions were prepared at 10 mM in DMSO and were diluted to the final working concentration in recording solution before use.

### Delayed alternation task

Rats were trained on the delayed alternation test of spatial working memory in a T-shaped maze. They were first adapted to handling and to eating treats (highly palatable miniature chocolate chips) on the maze prior to cognitive training. In this task, the rat was placed in the start box at the bottom of the ‘T’. When the gate was lifted, the rat proceeded down the stem of the maze to the choice point. On the first trial, rats were rewarded for entering either arm, but for each subsequent trial, were rewarded only if they chose the arm that they had not visited in the previous trial. Between trials, they were picked up and returned to the start box for a prescribed delay period. The choice point was cleaned with alcohol between each trial to remove olfactory clues (scent trails) often used by rodents to mark their previous locations. Successful performance of this task requires many operations carried out by the PFC: the rats must update and maintain the spatial information over the delay period for each trial, resist the distraction of being picked up and carried to the start box, and use response inhibition to overcome the tendency to repeat a rewarded action. The delay period was adjusted for each rat, such that they were performing at a stable baseline of 60–80% before drug treatment, thus leaving room for either impairment or improvement in performance. Rats were tested by experimenters who were highly familiar with the baseline behaviors of each individual animal, but blind to drug treatment conditions. Rats were observed for any potential differences in normative behavior (e.g. grooming, distracted sniffing), physical appearance, or physiological functioning (e.g. defecation/urination).

### Surgery and drug infusions into mPFC

Following training on the delayed alternation task, rats underwent aseptic stereotaxic surgery, under ketamine+xylazine anesthesia with metacam analgesic pretreatment, to implant cannula aimed just above the prelimbic mPFC (AP: +3.2 mm; ML: ±0.75 mm; DV: -4.2 mm). Details of the surgical procedure and post-surgical treatment can be found in (Gamo et al., 2015). After rats had fully recovered from the surgery, they were adapted to the infusion procedure to minimize stress. Once stable baseline performance was re-established, rats received intra-PFC infusions of vehicle vs. drug (0.5μl per side) over 5 minutes; performance was assessed 10 min after the infusion by a researcher unaware of the drug treatment conditions. There was at least a one-week washout between drug infusions.

### Analysis

Normality and variance similarity were measured by GraphPad Prism before we applied any parametric tests. Two-tailed Student’s t test was used for single comparisons between two groups. Other data were analyzed using one-way or two-way ANOVA with Bonferroni correction. All data were expressed as mean ± SEM, with statistical significance determined at p values < 0.05. In details, *Indicates *p <* 0.05; \*\**p <* 0.01; \*\*\**p <* 0.001; \*\*\*\**p <* 0.0001 in all figures.

## Figure Legends

**Figure 1S. Immunolabeling of Slack-B channels in Pyramidal neurons in layer II-III of mouse mPFC.** Note the reactive pyramidal somata (arrow) in layer II/III of mPFC slice from WT mice) and the lack of labelling of Slack-B protein in Slick/Slack Knockout mice.

