## Supplementary figures and images for "Interaction Between HCN and Slack Channels Regulates mPFC Pyramidal Cell Excitability and Working Memory"

### Supplemental Figure 1

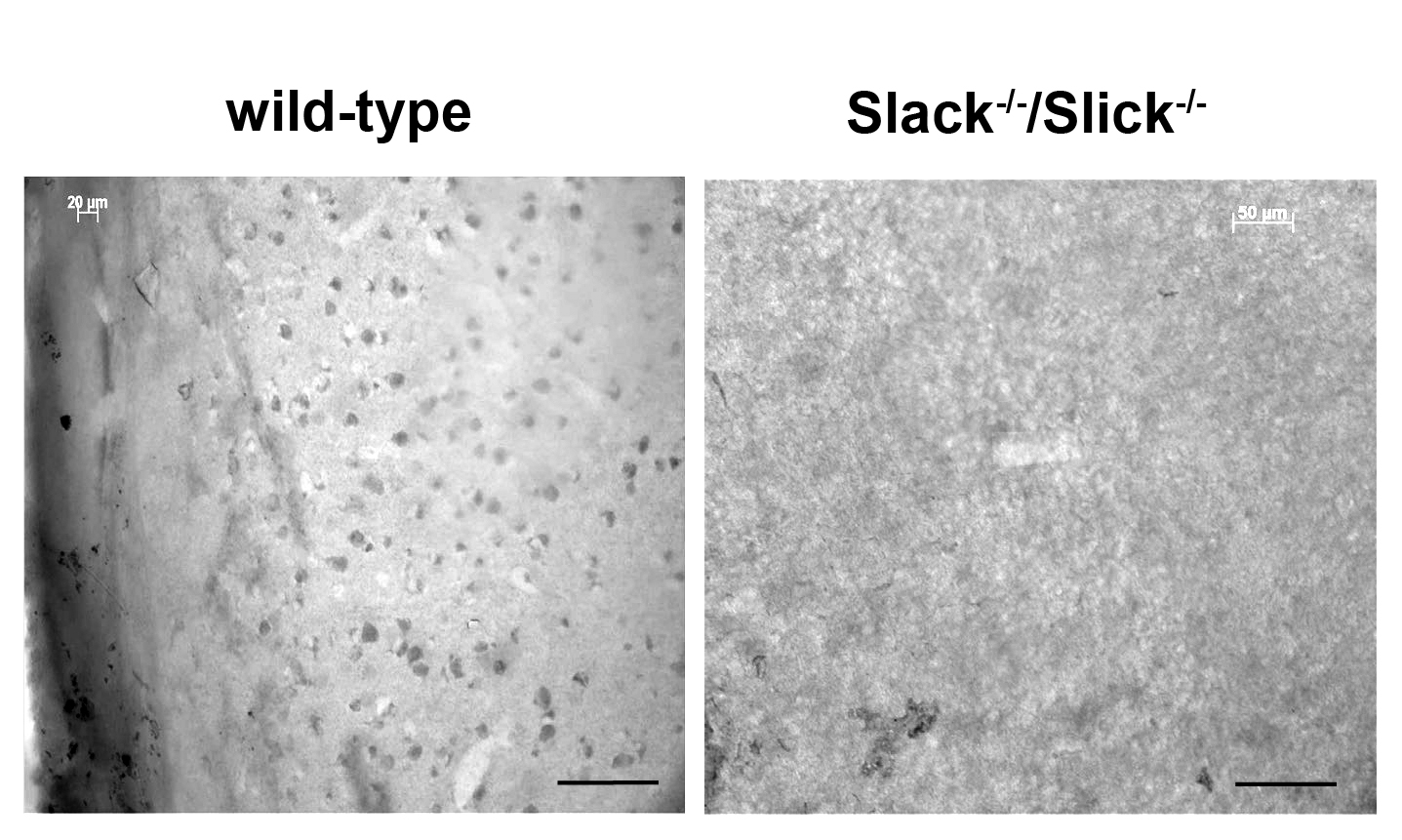
